# Context influences how individuals with misophonia respond to sounds

**DOI:** 10.1101/2020.09.12.292391

**Authors:** M. Edelstein, B. Monk, V.S. Ramachandran, R. Rouw

## Abstract

Misophonia is a newly researched condition in which specific sounds cause an intense, aversive response in individuals, characterized by negative emotions and autonomic arousal. Although virtually any sound can become a misophonic “trigger,” the most common sounds appear to be bodily sounds related to chewing and eating as well as other repetitive sounds. An intriguing aspect of misophonia is the fact that many misophonic individuals report that they are triggered more, or even only, by sounds produced by specific individuals, and less, or not at all, by sounds produced by animals (although there are always exceptions).

In general, anecdotal evidence suggests that misophonic triggers involve a combination of sound stimuli and contextual cues. The aversive stimulus is more than just a sound and can be thought of as a Gestalt of features which includes sound as a necessary component as well as additional contextual information. In this study, we explore how contextual information influences misophonic responses to human chewing, as well as sonically similar sounds produced by non-human sources. The current study revealed that the exact same sound can be perceived as being much more or less aversive depending on the contextual information presented alongside the auditory information. The results of this study provide a foundation for potential cognitive based therapies.

## INTRODUCTION

Misophonia is a newly researched condition in which specific sounds evoke an intensely aversive reaction in sufferers. Misophonia was first described by Jastreboff and Jastreboff (2001) nearly two decades ago but has only recently become a topic of interest to researchers in scientific and clinical communities. Sounds that evoke an intensely aversive reaction in individuals with misophonia are known as “triggers.” When exposed to these trigger sounds, individuals with misophonia experience a variety of physiological and negative emotional responses, resembling a fight-or-flight response (Edelstein et al., 2013; Brout et al., 2018; Kumar et al., 2014). At its most severe, misophonia can be so debilitating that it will often dictate the lives of those who suffer from it, causing people to go to great lengths just to avoid being exposed to certain sounds. Misophonic trigger sounds are frequently sounds that are not regarded as traditionally aversive to most individuals (although they may be considered annoying), and instead are commonly found to be human bodily noises (such as chewing, lip smacking, breathing or sniffing), or other repetitive sounds (such as tapping or pen clicking) (Schröder et al., 2013, Edelstein et al., 2013). While certain trigger sounds (such as chewing and mouthy sounds) appear to be far more common than others, it is important to note that each individual with misophonia possesses their own unique set of trigger sounds and that seemingly any sound has the potential to become a trigger.

When exposed to trigger sounds, misophonic individuals report experiencing intense feelings of anger, anxiety, disgust or rage (Schröder et al., 2013) in addition to a variety of physical sensations such as increased heart rate, tensing of muscles or perceived pressure building up in the body (Edelstein et al., 2013). It has been shown that, in response to auditory stimuli, including trigger sounds, misophonic individuals experience larger physiological responses (SCR and heart rate) indicative of autonomic nervous system arousal, than matched control participants (Edelstein et al., 2013; Kumar et al., 2017).

To date, only two published studies have explored the neural correlates associated with misophonia. Schröder et al. (2014) utilized electroencephalography (EEG) to measure auditory event related potentials (ERPs) in misophonic and control participants during an oddball task. They found that in response to oddball tones, misophonic but not control participants exhibited a decreased mean peak amplitude of the auditory N1 component, which is a component associated with early attention and detecting sudden changes in sensory information. As a decreased N1 component has been observed in individuals with a number of psychiatric conditions, the authors suggest that it could be interpreted as a marker of pathology and that misophonic individuals may be experiencing basic deficits in auditory processing. A groundbreaking study by Kumar et al. (2017) utilized neuroimaging techniques to highlight structural as well as functional neurological differences in those with and without misophonia. Findings revealed that in response to trigger sounds, misophonic participants showed increased activation in the bilateral anterior insular cortex (AIC) as well as increased functional connectivity between the AIC and regions of the brain associated with processing and regulating emotions. As the AIC is thought to be involved in the detection of important, salient stimuli, the increased activation found in misophonic individuals in response to trigger sounds suggests that these sounds are processed as being highly salient. In terms of structural differences, misophonic but not control participants were found to have increased myelination in the ventromedial prefrontal cortex (vmPFC), a region of the brain also involved in regulating emotions.

The prevalence of misophonia in the general population is not well understood yet. In a sample of 483 undergraduate students from a North American university, Wu et al. (2014) found that 20% reported experiencing symptoms of misophonia that were considered clinically significant. Additionally, a study by Zhou et al. (2017) which investigated the prevalence of misophonia in 415 students at two Chinese universities, found that while 16.6% reported clinically significant symptoms, only 6% were classified as experiencing significant levels of impairment. While these studies have made important contributions to our early understanding of misophonia, additional large-scale studies that sample a variety of populations are needed in order to gain an accurate sense of the true prevalence of the condition.

Misophonia has been found to be comorbid with conditions such as obsessive compulsive disorder (OCD) (Schröder et al., 2013; Wu et al., 2014; Ferreira et al., 2013), post-traumatic stress disorder (PTSD) (Rouw & Erfanian, 2017), depression (Wu et al., 2014), generalized anxiety disorder (Ferreira et al., 2013), ADHD (Rouw & Erfanian, 2017), Tourette’s syndrome (Neal & Cavanna, 2013), eating disorders (Kluckow et al., 2014) as well as tinnitus and hyperacusis (Jastreboff & Jastreboff, 2014). However, a significant number of individuals with misophonia report that they do not suffer from any additional conditions (Rouw & Erfanian, 2017). More research in this area is needed as there is currently no demonstrable evidence that a relationship exists between misophonia and other conditions (Potgieter et al., 2019).

A number of potential treatments for misophonia have been explored, including cognitive behavioral therapy (CBT) (Schröder et al., 2017; Bernstein et al., 2013; McGuire et al., 2015), tinnitus retraining therapy (TRT) (Jastreboff & Jastreboff, 2014), counterconditioning (Dozier, 2015), mindfulness and acceptance based approaches (Schneider & Arch, 2017) and pharmacological treatment (Vidal et al., 2017; Tunç et al., 2017). However, in addition to varying levels of effectiveness, there looms a significant problem in that these proposed treatments for misophonia are extremely preliminary and have not yet been validated through rigorous scientific testing (Potgieter et al., 2019).

In the last five years, misophonia has often been compared with another emerging sensory phenomenon called the autonomous sensory meridian response (ASMR) in which individuals experience pleasant tingling sensations (usually centralized around the scalp and neck) and feelings of relaxation in response to specific auditory and visual stimuli (Barratt & Davis, 2015; Janik McErlean & Banissy, 2018; Cash et al., 2018). ASMR inducing sounds (also termed “triggers”) often include whispering, quiet repetitive noises, crinkling, crisp sounds and sounds indicative of receiving personal attention. Interestingly, many ASMR triggers share striking similarities with misophonic triggers. Additionally, nearly half of the 300 misophonic participants in a study conducted by Rouw & Erfanian (2017) reported experiencing ASMR to certain sounds, suggesting a potential overlap between ASMR and misophonia that should undoubtedly be investigated further.

Despite a growing interest in misophonia in recent years, there still remains a marked lack of empirical research studies investigating the condition. The current study investigates an intriguing characteristic of misophonia reported by Edelstein et al. (2013) that may have the potential to inform future therapies. Namely, many sufferers have reported that sounds produced by certain individuals (typically family members and friends) are particularly aversive, while the same type of sound produced by another individual or a stranger may evoke less of a negative response or none at all. Also, self-produced trigger sounds rarely appear to evoke an aversive response in misophonic individuals. Given that an individual’s misophonia often appears to be localized around specific individuals, it seems like the misophonic response could be context sensitive. It has also been reported that the sounds of animals or babies are typically not found to be as aversive as similar sounding trigger sounds produced by adult humans. Although there are always exceptions, based on the aforementioned reports, it appears that an aversive stimulus often involves a highly nuanced formulation of sound and context, suggesting that a misophonic trigger is more than just a sound and instead, a Gestalt of features which includes sound (real or anticipated) as a necessary component. The idea that any singular feature of an aversive stimulus does not necessarily produce aversion on its own, is very interesting and warrants further exploration for both understanding misophonia on a fundamental level, and for its potential for clinically informative results.

Through the use of self-reported aversiveness ratings, we assessed participant aversion to a variety of classic trigger sounds in the presence and absence of contextual information. Clips of common trigger sounds (crunchy/wet human eating sounds) as well as sounds that highly resembled trigger sounds (crunchy/wet animal eating sounds and various crunchy/wet non eating sounds) were presented to self-identified misophonic and age/gender matched control participants in three experimental blocks. In each of the three experimental blocks, the type of contextual information accompanying each sound differed slightly. In block 1, participants were presented with only the audio of the sounds, and not given any feedback about what they were listening to. In block 2, participants were also presented with only the audio of the sounds, but prior to each sound, received a short text description about what they were potentially listening to. However, participants were informed that this description was not always correct and it was up to them to decide if the description matched the sound presented. In block 3, participants were presented with both the audio and video of each sound, which ultimately revealed the identity of each sound they had been listening to.

By utilizing deliberately ambiguous sounds and manipulating the type of contextual information provided about said sounds, our intention was to influence what participants believed they were listening to, to the extent where they may be convinced that certain trigger sounds were actually non-trigger sounds and certain non-trigger sounds were actually trigger sounds, and observe if their beliefs influenced their reactions. We hypothesized that misophonic individuals (but not controls) would find sounds that they perceived to be human eating sounds (regardless of whether they actually were or not) to be significantly more aversive than sounds that they perceived to be animal eating and non eating sounds. If successful, this study would demonstrate that contextual information that an individual associates with a sound can significantly influence their response to that sound, providing empirical evidence for the idea that the physical properties of a trigger sound are not the only factors driving the misophonic response.

## METHODS

### Participants

Twenty self-identified misophonic participants (5 males and 15 females; mean age = 30.4 years; range = 20-58) and twenty age and gender matched control participants (5 males and 15 females; mean age = 31.24 years; range = 20-58) were recruited from the student population at the University of California, San Diego and the greater San Diego area. All participants reported normal vision and hearing and signed a consent form approved by the UCSD Human Research Protections Program prior to participating. Participants were reimbursed with either UCSD course credit or at a rate of $10/hour. The entire lab session lasted for approximately 2 hours.

### Questionnaires

Control participants filled out a short demographic form that also assessed any prior knowledge of misophonia and sought to determine whether they may suffer from the condition unknowingly. No control participants were found to experience misophonic symptoms. Self-identified misophonic participants were given a demographic form as well as several commonly used misophonia questionnaires that assessed their experiences with the condition and gauged the severity of their symptoms. The questionnaires included were the Amsterdam Misophonia Scale (A-MISO-S), which measures the severity of the symptoms and intensity of responses associated with a participant’s misophonia, the Misophonia Activation Scale (MAS-1) which characterizes eleven levels (0-10) of misophonia severity, and the Misophonia Assessment Questionnaire (MAQ) which assesses how frequently participants experience negative effects and disturbances associated with misophonia.

Misophonic participants scored an average of 11.7 (range = 7-24) points out of a maximum of 24 (most severe) points on the A-MISO-S and an average of 28 (range: 10-63) points out of a maximum of 63 points (most severe) on the MAQ. Of the eleven levels of misophonia severity detailed in the MAS-1 (0-10), the average level amongst participants was found to be 5.475 (range = 3.5-9).

### Experimental Setup

As a general overview, each participant took part in a session that consisted of 3 experimental blocks. Although it differed slightly from block-to-block, the general structure of a block was as follows: participants were seated 20 inches away from a computer screen and wore a pair of Sennheiser headphones. Through the use of MATLAB R2014B, visual stimuli were presented on the computer screen and auditory stimuli were presented through the headphones at 50% of the computer’s volume. An individual trial consisted of a 5 second (pre-stimulus period) followed by a 15 second clip (stimulus period), and finally a 10 second intertrial interval (ITI). During the ITI, participants were instructed to verbally make an aversiveness rating about the clip they were just presented with on a 1-10 scale. Each block contained 36 clips, with each clip falling into one of three sound categories (Fig. 1).

**Figure 1.**
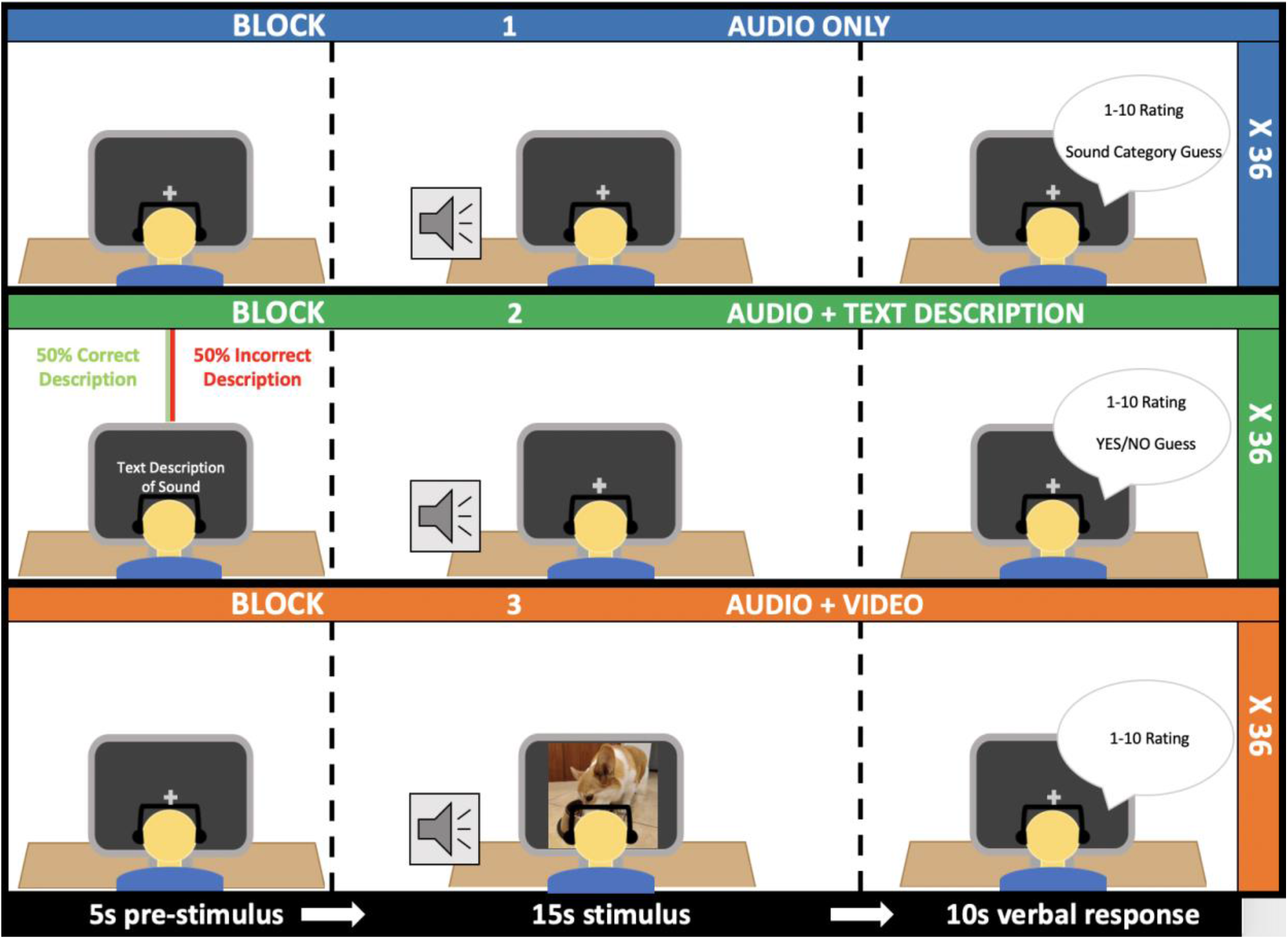
Experimental setup and procedure.

Participants were informed beforehand that an aversiveness rating of “1” signified very little to no discomfort while a rating of “10” signified extreme discomfort and possibly a strong desire to leave the room should the sound continue. Each aversiveness rating was recorded by the experimenter. Between blocks, participants were instructed to take a short break.

### Stimuli

Thirty-six, 15-second video clips were used in the study. All clips were either found on Youtube or created in the lab. Each clip was placed into one of three sound categories: human eating, animal eating or non eating, with 12 clips in each category. Clips were selected based on the criteria that they either were or highly resembled classic misophonic trigger sounds (most were crunchy or wet sounding in nature). Audio (sound only) and audio-visual (sound + video) versions of each clip were created.

Clips were selected based on results from a pilot study involving 21 participants that was conducted in the summer of 2016. The purpose of this pilot study was to identify a set of classic misophonic sounds that could plausibly be interpreted as belonging to more than one of the aforementioned sound categories (when presented with only audio and no visuals). The most categorically ambiguous clips were then selected to be used as stimuli in the current study.

### Experimental Blocks

For each block 1 trial, participants were presented with a 5 second pre-stimulus period followed by a 15 second audio only clip, and then a 10 second ITI during which they made their aversiveness rating on a 1-10 scale. In addition to their aversiveness rating, participants were also instructed to make a guess as to what they thought the sound source of each clip was (based on the aforementioned 3 sound categories) during this ITI. The sounds in block 1 were presented in a randomized order for every participant (Fig. 1).

For each block 2 trial, participants were presented with a 5 second pre-stimulus period which included 1 second of blank, black screen, followed by 3 seconds of descriptive text, followed by 1 second of blank, black screen. Next came a 15 second audio only clip and then the 10 second ITI during which participants made their aversiveness rating on a 1-10 scale. Half of the time the text presented during the pre-stimulus period was a correct description of the sound that would play immediately after it and half of the time it was an incorrect description (Fig. 1). When it was incorrect, the text was a randomly selected description from one of the other two sound categories that the sound from that trial did not fall under. An incorrect description was never from the same category as the sound presented. For each trial in block 2, participants responded with a “yes” or “no” as to whether or not the text description they received sounded like the sound they were presented with. Participants were instructed to make this judgment based on general sound category and not the specifics of each description. The ordering of both the textual descriptions and sounds were preselected for each participant and counterbalanced. There were 6 possible sound and 6 possible text pseudo-random orderings. Each misophonic participant was matched with a control participant who received the same sound and text ordering in block 2 as they did.

For each block 3 trial, participants were presented with a 5 second pre-stimulus period followed by a 15 second video clip (audio and video) and then a 10 second ITI during which participants made their aversiveness rating on a 1-10 scale. Video clips in block 3 were presented in a randomized order for each participant (Fig. 1).

## RESULTS

### Within Blocks Results

#### Block 1: Audio Only | All Trials

As expected, we found that overall, aversiveness ratings given by misophonics (M = 4.92) were significantly higher than ratings given by controls (M = 1.97) [F(1,38) = 49.764, p < .001]. There was also an observed within subject factor effect of Sound Category [F(2,76) = 25.719, p < .001], where aversion to sounds from the human eating category (M = 4.01) was significantly higher than aversion to sounds from the animal eating (M = 3.18) and non eating (M = 3.15) categories, across groups (Fig. 2A).

**Figure 2.**
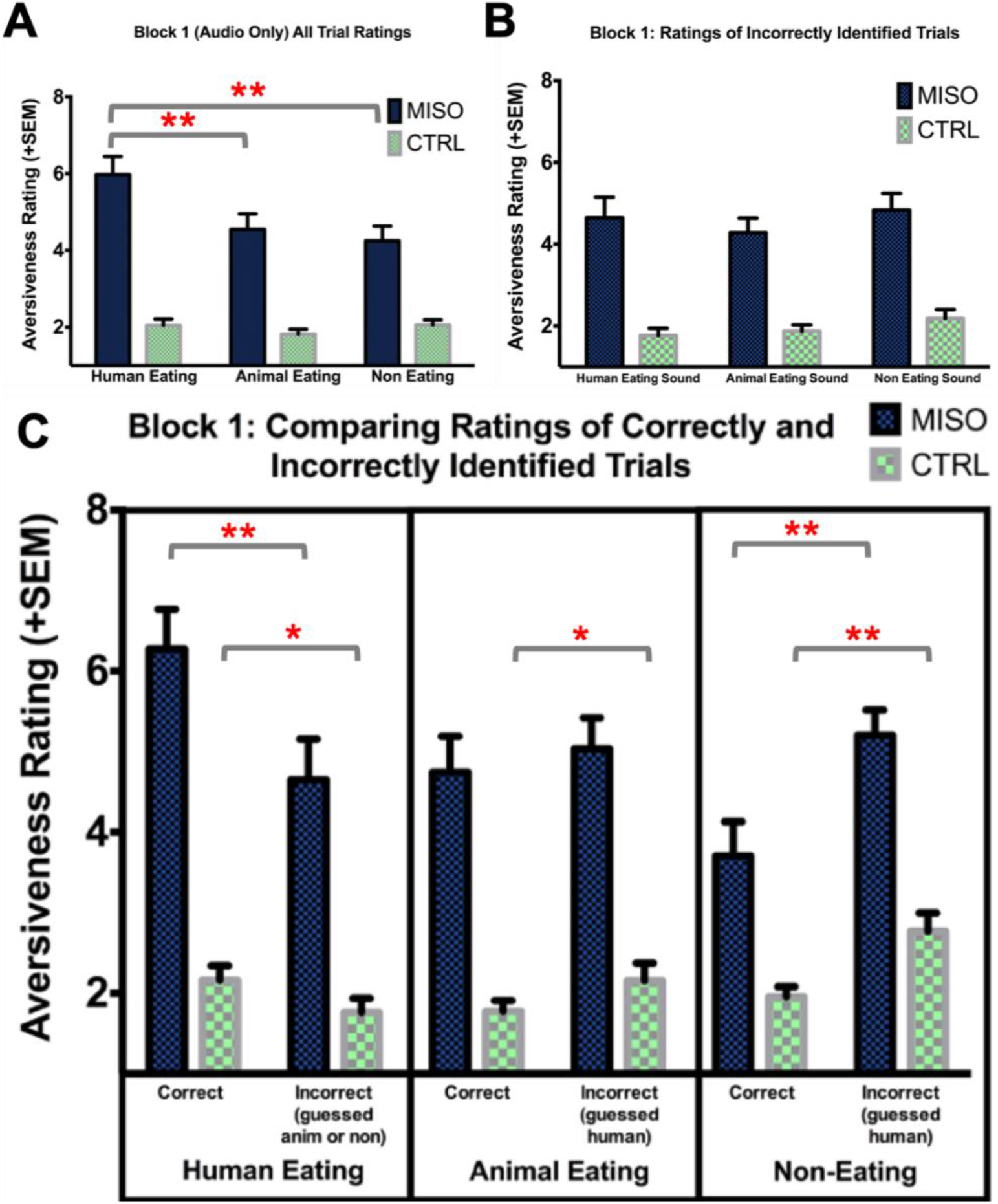
Block 1 aversiveness ratings. A) Average aversiveness ratings of human eating, animal eating and non eating sounds for misophonic and control participants in block 1, regardless of if the sound category was correctly identified. B) Average aversiveness ratings of incorrectly identified human eating, animal eating and non eating sounds for misophonic and control participants in block 1. C) Average aversiveness ratings of correctly and specific incorrectly identified human eating, animal eating and non eating sounds of misophonic and control participants in block 1. Error bars represent the standard error of the mean. * p < 0.05, ** p < 0.01

This observed main effect of Sound Category was driven by a significant interaction between Group and Sound Category [F(2,76) = 21.406, p < .001], where misophonic participants rated human eating sounds as particularly aversive compared with animal eating and non eating sounds (M = 5.98, 4.55, 4.25 respectively for misophonics; M = 2.05, 1.82, 2.06 respectively for controls). As a follow up to the interaction, paired t-tests indicated that misophonic participants rated human eating sounds as significantly more aversive than both animal eating [t(19) = 6.36, p < .001] and non eating [t(19) = 5.27, p < .001] sounds (there was no statistical difference between animal eating vs non eating sounds [t(19) = 1.563, p = .135]) (Fig. 2A).

Figure 4 depicts average misophonic and control ratings of each stimulus from block 1 in the form of a scatterplot, illustrating the finding that misophonic participants found all stimuli to be more aversive than controls did as well as showing which sounds were found to be most aversive.

#### Block 1: Audio Only | Correct Trials

For ratings of trials where participants correctly identified the sound category, we found significant main effects of Group [F(1,38) = 48.109, p < .001], with the misophonic group rating sounds as significantly more aversive overall (M = 4.91) than the control group (M = 1.97), and Sound Category [F(2,76) = 28.552, p < .001], with human eating sounds rated as more aversive (M = 4.22) than animal eating (M = 3.26) and non eating sounds (M = 2.83) across groups. A significant interaction between Group and Sound Category [F(2,76) = 19.801, p < .001] was also observed, with human eating sounds rated as particularly aversive by the misophonic participants (M = 6.28), compared with animal eating (M = 4.75) and non eating sounds (M = 3.71) (Fig. 2C).

Paired t-tests confirmed that misophonics rated human eating sounds as significantly more aversive than animal eating sounds [t(19) = 5.09, p < .001] and non eating sounds [t(19) = 5.60, p < .001]. Interestingly, misophonics also rated animal eating sounds as significantly more aversive than non eating sounds [t(19) = 3.79, p = .001]. Controls were found to rate human eating sounds as significantly more aversive than animal eating sounds, [t(19) = 3.13, p = .006], but not non eating sounds [t(19) = 1.50, p = .149]. Controls also found non eating sounds to be marginally more aversive than animal eating sounds [t(19) = 1.826, p = .084].

#### Block 1: Audio Only | Incorrect Trials

For ratings of trials where the participant incorrectly identified the sound category, we found a significant main effect of Group [F(1,38) = 42.685, p < .001], with the misophonic group rating sounds as significantly more aversive overall (M = 4.59) than the control group (M = 1.94), a marginally significant main effect of Sound Category [F(2,76) = 2.557, p = .084], (human eating sounds (M = 3.21), animal eating (M = 3.08), non eating sounds (M = 3.51) across groups), but no significant interaction between Group and Sound Category [F(2,76) = .593, p = .475], with human eating sounds not rated as particularly aversive by the misophonic participants (M = 4.65) when compared with animal eating (M = 4.28) and non eating sounds (M = 4.84) (Fig. 2B).

Paired t-tests indicated that misophonics did not demonstrate any significant difference in aversiveness ratings between incorrectly identified human eating sounds and incorrectly identified animal eating sounds [t(19) = .938, p = .36] or incorrectly identified non eating sounds [t(19) = −.59, p = .562]. There was also no significant difference between the ratings of incorrectly identified animal eating sounds and incorrectly identified non eating sounds [t(19) = −1.67, p = .112] for misophonics. Controls did not demonstrate any significant difference in aversiveness ratings between incorrectly identified human eating sounds and incorrectly identified animal eating sounds [t(19) = −.892, p = .384] or incorrectly identified non eating sounds [t(19) = −1.80, p = .088]. There was also no significant difference between the ratings of incorrectly identified animal eating sounds and incorrectly identified non eating sounds [t(19) = −1.582, p = .130] for controls.

#### Block 1: Audio Only | Trial Comparisons

Additionally, the ratings of trials where the participant incorrectly identified the sound category were compared to trials where they correctly identified the sound category. Specifically, we were interested in comparing 1) ratings of trials where human eating sounds were misidentified as either animal eating sounds or non eating sounds to ratings trials where human eating sounds were correctly identified as human eating sounds, 2) ratings of trials where animal eating sounds were misidentified as human eating sounds to ratings of trials where animal eating sounds were correctly identified as animal eating sounds and 3) ratings of trials where non eating sounds were misidentified as human eating sounds to ratings of trials where non eating sounds were correctly identified as non eating sounds.

Results showed that misophonics rated human eating sounds incorrectly identified as animal eating or non eating sounds (M = 4.64, SD = 2.26) as significantly less aversive than human eating sounds correctly identified as human eating sounds (M = 6.28, SD = 2.18), [t(19) = −4.46, p < .001]. The same pattern was present for controls [t(19) = −2.2, p = .04]. Although there was no significant difference between misophonic ratings of animal eating sounds incorrectly identified as human eating sounds (M = 5.04, SD = 1.7) and animal eating sounds correctly identified as animal eating sounds (M = 4.75, SD = 1.97), [t(19) = −.865, p = .398]s, controls did show a significant difference in ratings between these two groups of trials, [t(19) = −2.25, p = .036]. Lastly, misophonics rated non eating sounds incorrectly identified as human eating sounds (M = 5.2, SD = 1.41) as significantly more aversive than non eating sounds correctly identified as non eating sounds (M = 3.7, SD = 1.91), [t(19) = 3.08, p = .006]. Controls also exhibited the same pattern of results for non eating sounds [t(19) = 3.3, p = .004] (Fig. 2C).

#### Block 1: Audio Only: Stimulus Classification | Category Guess Propensity & Accuracy

We also investigated the level of accuracy for sound category identification (percentage of trials correct) with factors of Group (misophonics, controls) and Sound Category (human eating, animal eating, non eating). No significant main effect of Group [F(1,38) = .615, p = .438] was observed, but there was a significant main effect of Sound Category [F(2,76) = 18.156, p < .001] as well as a marginally significant interaction between Group and Sound Category [F(2,76) = 3.353, p = .04]. This interaction brings about a few interesting findings. The first finding revealed that misophonic participants (M = 79.175%, SD = 16.120%) were significantly more accurate than controls (M = 67.075%, SD = 12.8293%) when identifying human eating sounds in particular [F(1,38) = 6.899, p = .012] but not when identifying animal eating sounds [F(1,38) = .034, p = .854] or non eating sounds [F(1,38) = 1.756, p = .193] (Fig. 3A).

**Figure 3.**
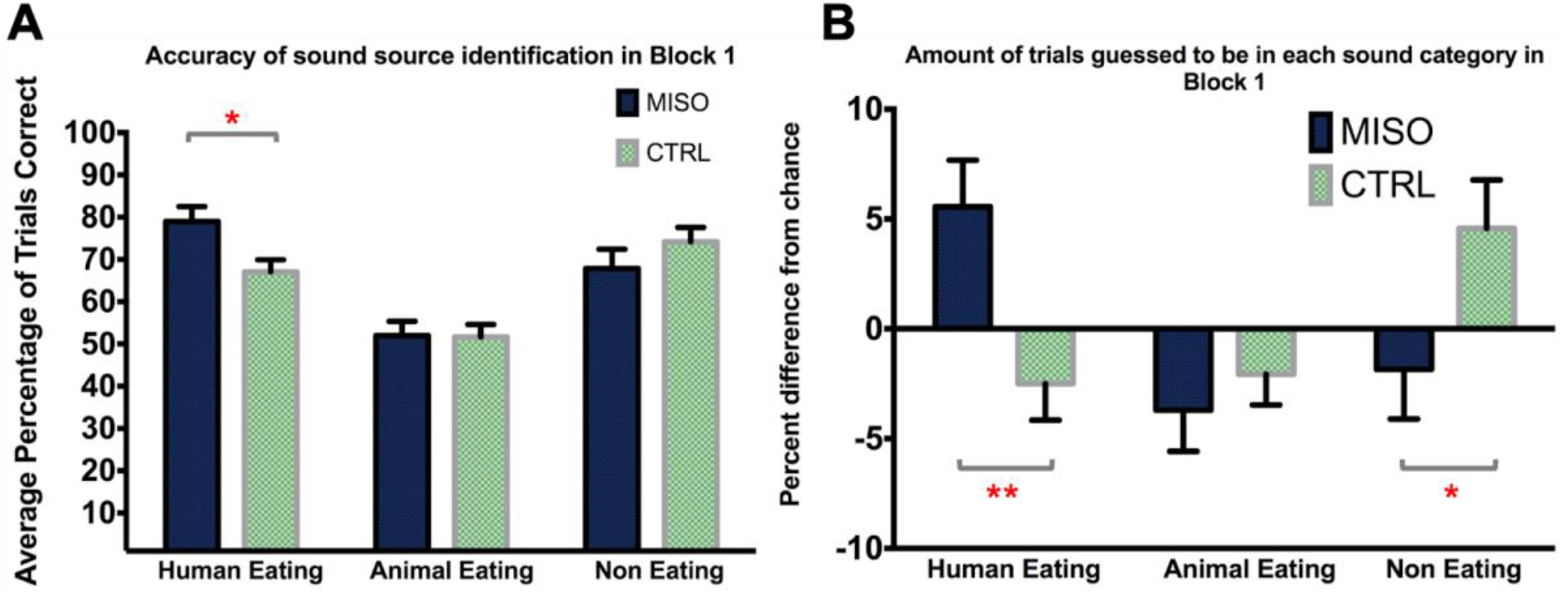
Sound category guess accuracy and propensity. A) Average percentage of correct human eating, animal eating and non eating trials of misophonic and control participants in block 1. B) Depicts how frequently trials (shown as percent difference from chance (33.33%)) were guessed by misophonic and control participants to be human eating, animal eating and non eating sounds in block 1. Error bars represent the standard error of the mean. * p < 0.05, ** p < 0.01

**Figure 4.**
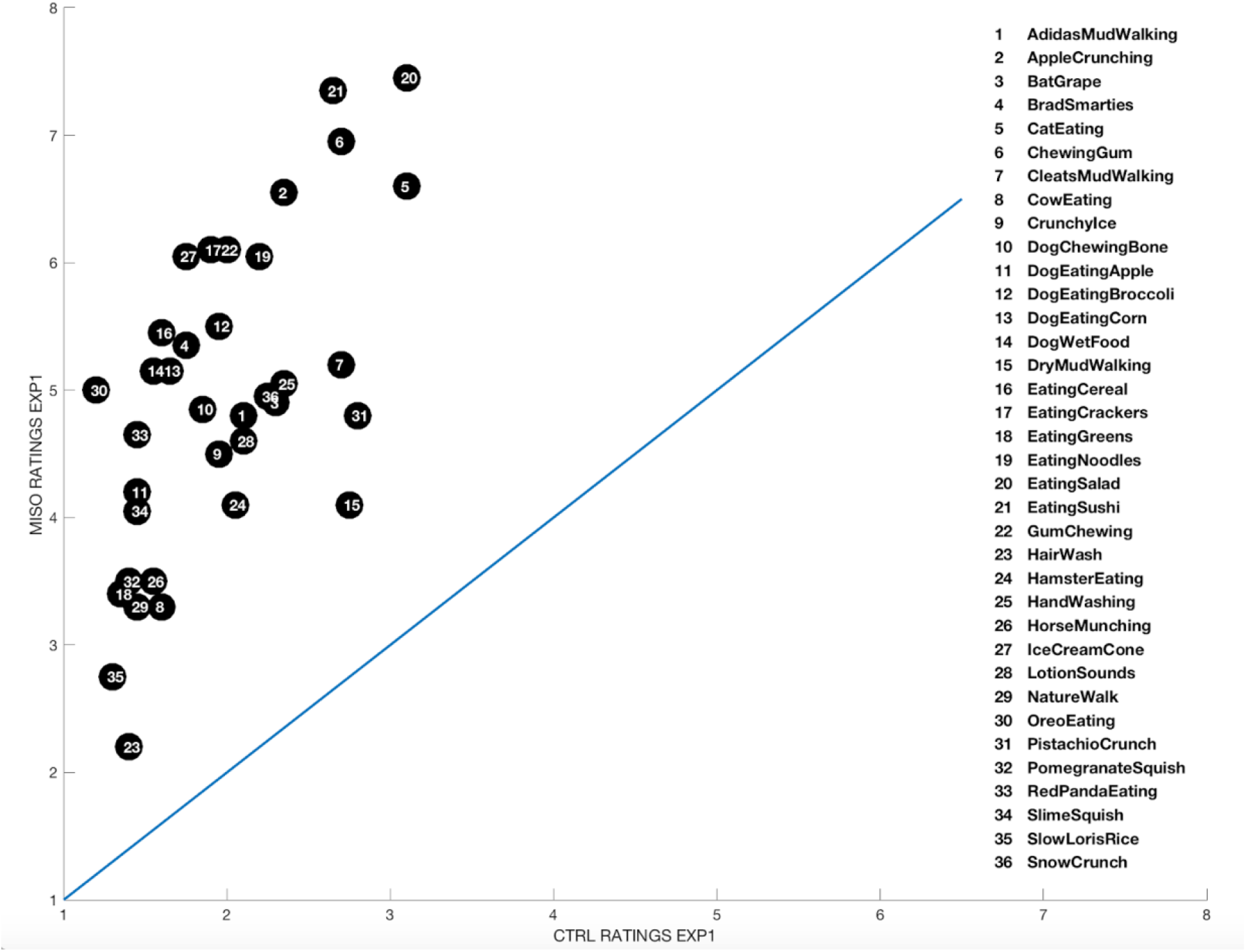
Scatterplot of misophonic and control ratings of all stimuli in block 1.

Paired t-tests indicated that misophonics were also significantly more accurate at identifying human eating sounds than animal eating sounds [t(19) = 5.361, p < .001], significantly more accurate at identifying non eating sounds than animal eating sounds [t(19) = 2.482, p = .023] and marginally more accurate at identifying human eating sounds than non eating sounds [t(19) = 1.885, p = .075]. Additionally, controls were significantly more accurate at identifying human eating sounds than animal eating sounds, [t(19) = 4.708, p < .001], significantly more accurate at identifying non eating sounds than animal eating sounds [t(19) = 4.162, p = .001] but not significantly more accurate at identifying non eating sounds than human eating sounds [t(19) = 1.256, p = .224].

In order to address the possibility that participants may have demonstrated a preference to make guesses within a specific sound category (which could influence their accuracy), the percentage of trials that were guessed to be in each sound category was investigated with factors of Group (misophonics, controls) and Sound Category (human eating, animal eating, non eating). Although no significant main effects of Group [F(1,38) = .580, p = .451] or Sound Category [F(2,76) = 1.9, p = .157] were observed, a significant interaction between Group and Sound Category [F(2,76) = 5.472, p = .006] was found. This interaction brings about a few interesting findings. The first finding revealed that misophonic participants were significantly more likely than controls to guess that a sound was a human eating sound [F(1,38) = 9.37, p = .004] but not significantly more likely than controls to guess that a sound was an animal eating sound [F(1,38) = .233, p = .632]. Interestingly, controls were significantly more likely than misophonics to guess that a sound was a non eating sound [F(1,38) = 5.395, p = .026] (Fig. 3B).

Paired t-tests indicated that misophonics were also significantly more likely to guess that a sound was a human eating sound as opposed to an animal eating sound [t(19) = 2.667, p = .015], or a non eating sound [t(19) = 2.16, p = .044]. No significant difference in guessing rate was found between animal eating and non eating sounds [t(19) = .164, p =.871]. Additionally, although controls were not significantly more likely to guess that a sound was a human eating sound as opposed to an animal eating sound [t(19) = .202, p = .842], they were marginally more likely to guess that a sound was a non eating sound as opposed to a human eating sound [t(19) = 1.936, p = .068], or an animal eating sound [t(19) = 2.011, p = .059].

#### Block 2: Audio + Text | Agree + Disagree Trials

First, we conducted a repeated measures mixed design ANOVA on factors of Group (misophonic, control) and Sound Category (human eating, animal eating, non eating) for ratings of all block 2 trials (regardless of whether the participant received an accurate (target) or false (foil) textual description and regardless of whether they got the trial right or wrong). Overall, we found significant main effects of Group [F(1,38) = 51.1, p < .001] and Sound Category [F(2,76) = 34.7, p < .001] as well as a significant interaction between Group and Sound Category [F(2,76) = 27.6, p < .001].

Follow up paired t-tests revealed that misophonics rated human eating sounds (M = 6.14, SD = 2.03) as significantly more aversive than both animal eating (M = 5.00, SD = 2.01), [t(19) = 5.87, p < .001] and non eating sounds (M = 4.45, SD = 1.8), [t(19) = 6.69, p <.001]. Additionally, misophonics rated animal eating sounds as significantly more aversive than non eating sounds [t(19) = 3.37, p = .003]. Controls demonstrated a similar pattern of results and rated human eating sounds (M = 2.1, SD = .79) as significantly more aversive than animal eating sounds (M = 1.87, SD = .6), [t(19) = 3.301, p = .004] but not non eating sounds (M = 2.05, SD = .719) [t(19) = .669, p = .512]. Additionally, controls rated non eating sounds as significantly more aversive than animal eating sounds [t(19) = 2.75, p = .013] (Fig. 5A).

**Figure 5.**
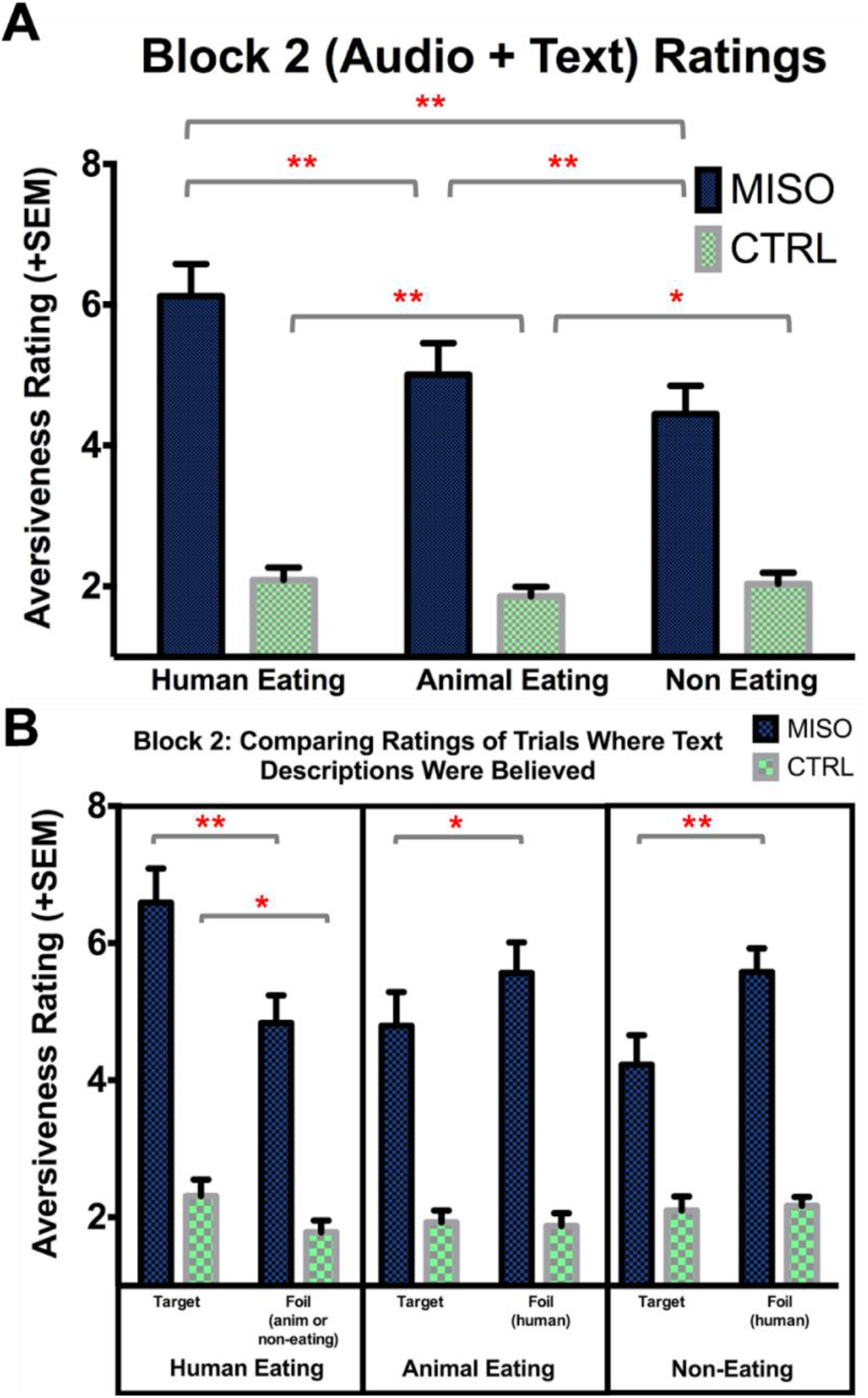
Block 2 aversiveness ratings. A) Average aversiveness ratings of human eating, animal eating and non eating sounds for misophonic and control participants in block 2, regardless of whether participants received an accurate (target) or false (foil) textual description and whether the sound category was correctly identified. B) Misophonic and control participants average aversiveness ratings of human eating, animal eating and non eating sounds that were either correctly identified as their target description (left) or incorrectly identified as specific foil descriptions (right) in block 2. Error bars represent the standard error of the mean. * p < 0.05, ** p < 0.01

#### Block 2: Audio + Text | Agree Trials

We compared the ratings of trials where participants incorrectly believed false text (foil) descriptions preceding the stimulus to trials where they correctly believed true text (target) descriptions preceding the stimulus. Specifically, we were interested in comparing 1) ratings of trials where human eating sounds were incorrectly believed to be either animal eating sounds or non eating sounds to ratings of trials where human eating sounds were correctly believed to be human eating sounds, 2) ratings of trials where animal eating sounds were incorrectly believed to be human eating sounds to ratings of trials where animal eating sounds were correctly believed to be animal eating sounds and 3) ratings of trials where non eating sounds were incorrectly believed to be human eating sounds to ratings of trials where non eating sounds were correctly believed to be non eating sounds.

Specifically, misophonics rated human eating sounds incorrectly believed to be animal eating or non eating sounds (M = 4.83, SD = 1.81) as significantly less aversive than human eating sounds correctly believed to be human eating sounds (M = 6.59, SD = 2.23), [t(19) = −4.344, p < .001]. Misophonics also rated animal eating sounds incorrectly believed to be human eating sounds (M = 5.56, SD = 2.00) as significantly more aversive than animal eating sounds correctly believed to be animal eating sounds (M = 4.79, SD = 2.21), [t(19) = 1.69, p = .05]. Additionally, misophonics rated non eating sounds incorrectly believed to be human eating sounds (M = 5.58, SD = 1.55) as significantly more aversive than non eating sounds correctly believed to be eating sounds (M = 4.23, SD = 1.92), [t(19) = 3.03, p = .0035] (Fig. 5B).

Controls did not rate animal eating sounds incorrectly believed to be human eating sounds as significantly more aversive than animal eating sounds correctly believed to be animal eating sounds [p >.05]. They also did not rate non eating sounds incorrectly believed to be human eating sounds as significantly more aversive than non eating sounds correctly believed to be eating sounds [p >.05]. However, controls did rate human eating sounds incorrectly believed to be animal eating or non eating sounds (M = 1.78, SD = .76) as significantly less aversive than human eating sounds correctly believed to be human eating sounds (M = 2.31, SD = 1.07), [t(19) = −2.13, p = .047] (Fig. 5B).

#### Block 3: Audio + Video Trials

We conducted a repeated measures mixed design ANOVA on factors of Group (misophonic, control) and Sound Category (human eating, animal eating, non eating) for ratings of block 3 trials. Overall, we found significant main effects of Group [F(1,38) = 46.822, p < .001] and Sound Category [F(2,76) = 51.879, p < .001] as well as a significant interaction between Group and Sound Category [F(2,76) = 21.081, p < .001].

Follow up paired t-tests revealed that misophonics rated human eating sounds (M = 6.97, SD = 2.01) as significantly more aversive than both animal eating (M = 3.99, SD = 2.28), [t(19) = 7.39, p < .001] and non eating sounds (M = 6.57, SD = 2.08), [t(19) = 6.69, p < .001]. Misophonics did not rate animal eating sounds as significantly more aversive than non eating sounds [t(19) = −.186, p = .854]. Controls rated human eating sounds (M = 2.44, SD = .87) as significantly more aversive than animal eating sounds (M = 1.63, SD = .46), [t(19) = 5.134, p < . 001] and non eating sounds (M = 1.94, SD = .63) [t(19) = 3.13, p = .006]. Additionally, controls rated non eating sounds as significantly more aversive than animal eating sounds [t(19) = 3.29, p = .004] (Fig. 6).

**Figure 6.**
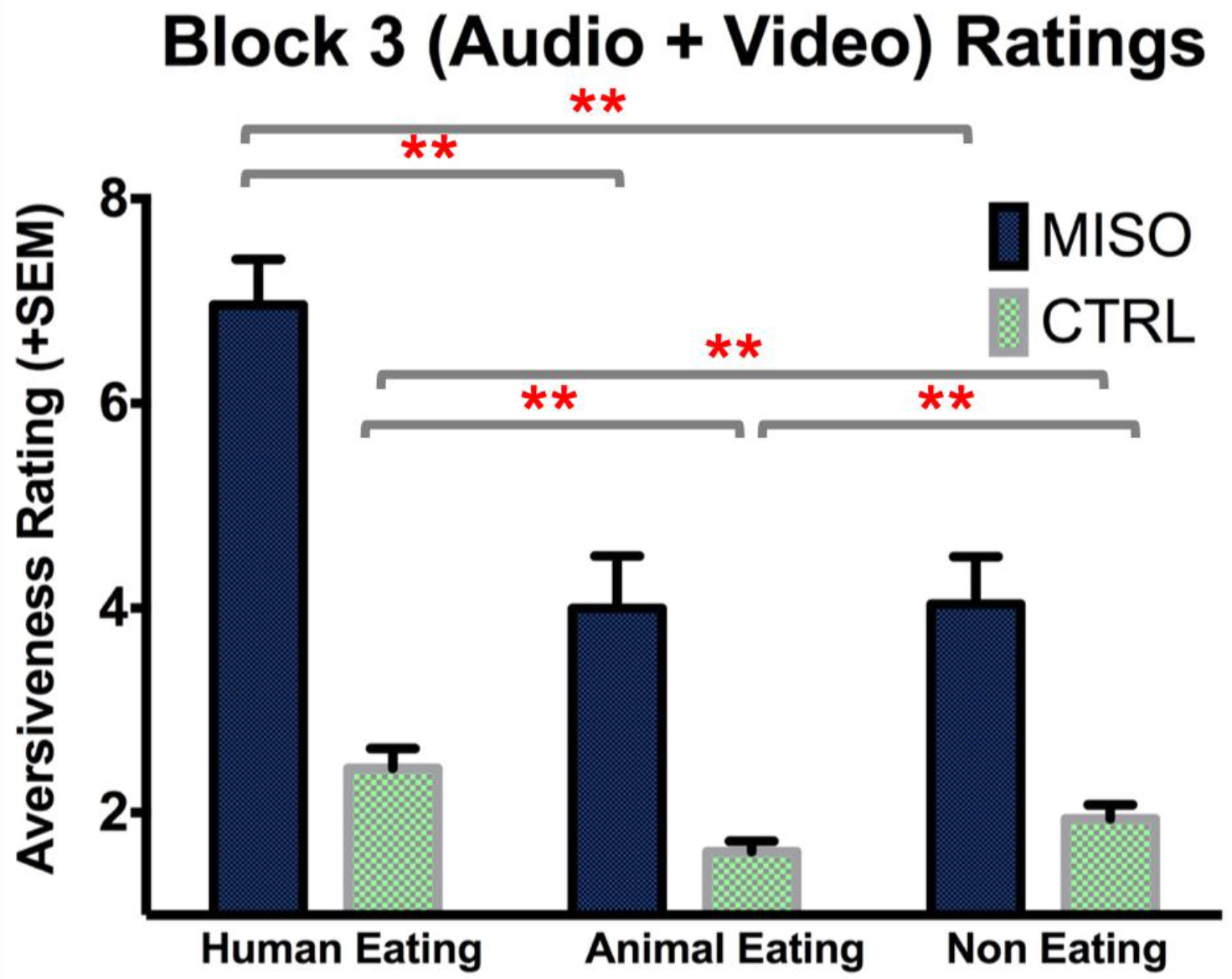
Block 3 aversiveness ratings. Average aversiveness ratings of human eating, animal eating and non eating sounds for misophonic and control participants in block 3. Error bars represent the standard error of the mean. * p < 0.05, ** p < 0.01

### Between Blocks Results

In addition to investigating how misophonic and control participants responded to human eating, animal eating and non eating sounds within the differing contexts of blocks 1, 2 and 3, we also examined how their responses to specific sounds changed across these blocks. In particular, we were interested in observing how their ratings changed between all three blocks for

1. human eating sounds that were correctly identified as human eating sounds in block 1 but were believed to be produced by nonhuman (animal eating and non eating sounds) sources in block 2.
2. nonhuman sounds (animal eating and non eating sounds) that were correctly identified as nonhuman sounds in block 1 but were believed to be human eating sounds in block 2.
3. human eating sounds that were correctly identified as human eating sounds in blocks 1 and 2.
4. nonhuman sounds (animal eating and non eating sounds) that were correctly identified as nonhuman sounds in blocks 1 and 2.

When considering human eating sounds that were correctly identified as human eating sounds in block 1 but were believed to be produced by nonhuman sources in block 2, we find that misophonics rated these sounds to be significantly more aversive in block 1 (M = 5.95, SD = 1.69) than when they encountered them again in block 2 (M = 5.0, SD = 1.58), [t(13) = 1.892, p = .04]. Misophonics additionally rated these sounds to be significantly more aversive in block 3 (M = 6.45, SD = 1.45) than block 2 (M = 5.0, SD = 1.58), [t(13) = 3.065, p = .0045], but no significant difference in ratings for these sounds was found between blocks 1 and 3, [p > .05]. Controls did not exhibit significant differences in ratings for these sounds between blocks 1 and 2 [p > .05], blocks 2 and 3 [p > .05] or blocks 1 and 3 [p > .05] (Fig. 7A).

**Figure 7.**
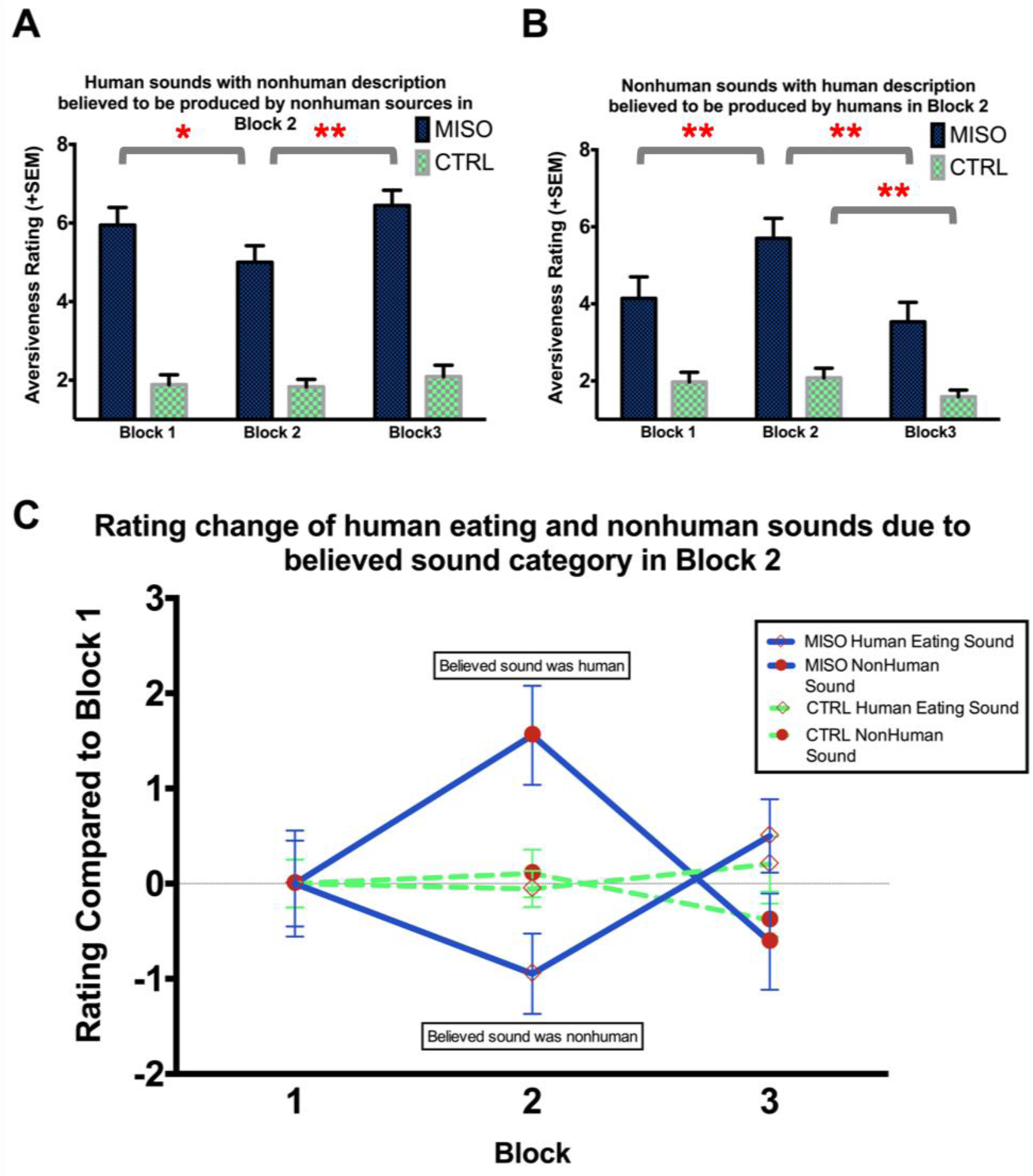
Rating change of human eating sounds and nonhuman sounds (believed to be the opposite type of sound in block 2) across blocks. A) Depicts the average misophonic and control aversiveness ratings of human eating sounds that were correctly identified as human eating sounds in block 1 but were believed to be produced by nonhuman sources in block 2. B) Depicts the average misophonic and control aversiveness ratings of nonhuman sounds that were correctly identified as nonhuman sounds in block 1 but were believed to be human eating sounds in block 2. C) Combines figures 7A and 7B into one graph but instead of displaying average aversiveness ratings, displays the average change in rating of each sound type in each block relative to block 1.

When considering nonhuman sounds (animal eating and non eating sounds) that were correctly identified as nonhuman sounds in block 1 but were believed to be human eating sounds in block 2, we find that misophonics rated these sounds to be significantly more aversive in block 2 (M = 5.7, SD = 2.32) than in block 1 (M = 4.14, SD = 2.5), [t(19) = 4.098, p < . 001]. Misophonics also rated these sounds as significantly more aversive in block 2 (M = 5.7, SD = 2.32) than in block 3 (M = 3.53, SD = 2.26), [t(19) = 4.875, p < .001], but no significant difference in ratings for these sounds was found between blocks 1 and 3, [p > .05]. Controls did not exhibit significant differences in ratings for these sounds between blocks 1 and 2 [p > .05] or blocks 1 and 3 [p > .05], but a significant difference between blocks 2 and 3 was observed, with these sounds being rated as significantly more aversive in block 2 (M = 2.08, SD = 1.03) than block 3 (M = 1.59, SD = .712), [t(16) = 3.125, p = .007] (Fig. 7B).

When comparing human eating sounds that misophonics correctly identified as human eating sounds in block 1 but were believed to be produced by nonhuman sources in block 2, with nonhuman sounds (animal eating and non eating sounds) that misophonics correctly identified as nonhuman sounds in block 1 but were believed to be human eating sounds in block 2, we find a significant main effect of the between subject factor of Sound Type [F(1,32) = 4.57, p = .04] and a significant interaction between Sound Type and the within subject factor Block [F(2,64) = 17.254, p < .001] (Fig. 7C).

When considering human eating sounds that were correctly identified as human eating sounds in blocks 1 and 2, we find that misophonics rated these sounds to be marginally more aversive in block 2 (M = 6.6, SD = 2.29) than when they encountered them in block 1 (M = 6.32, SD = 2.36), [t(19) = 1.48, p = .08], significantly more aversive in block 3 (M = 7.07, SD = 2.15) than block 2 (M = 6.6, SD = 2.29), [t(19) = 2.33, p = .015], and significantly more aversive in block 3 (M = 7.07, SD = 2.15) than block 1 (M = 6.32, SD = 2.36), [t(19) = 3.28, p = .002]. Controls did not exhibit significant differences in ratings for these sounds between blocks 1 and 2 [p > .05], blocks 2 and 3 [p > .05] or blocks 1 and 3 [p > .05] (Fig. 8A).

**Figure 8.**
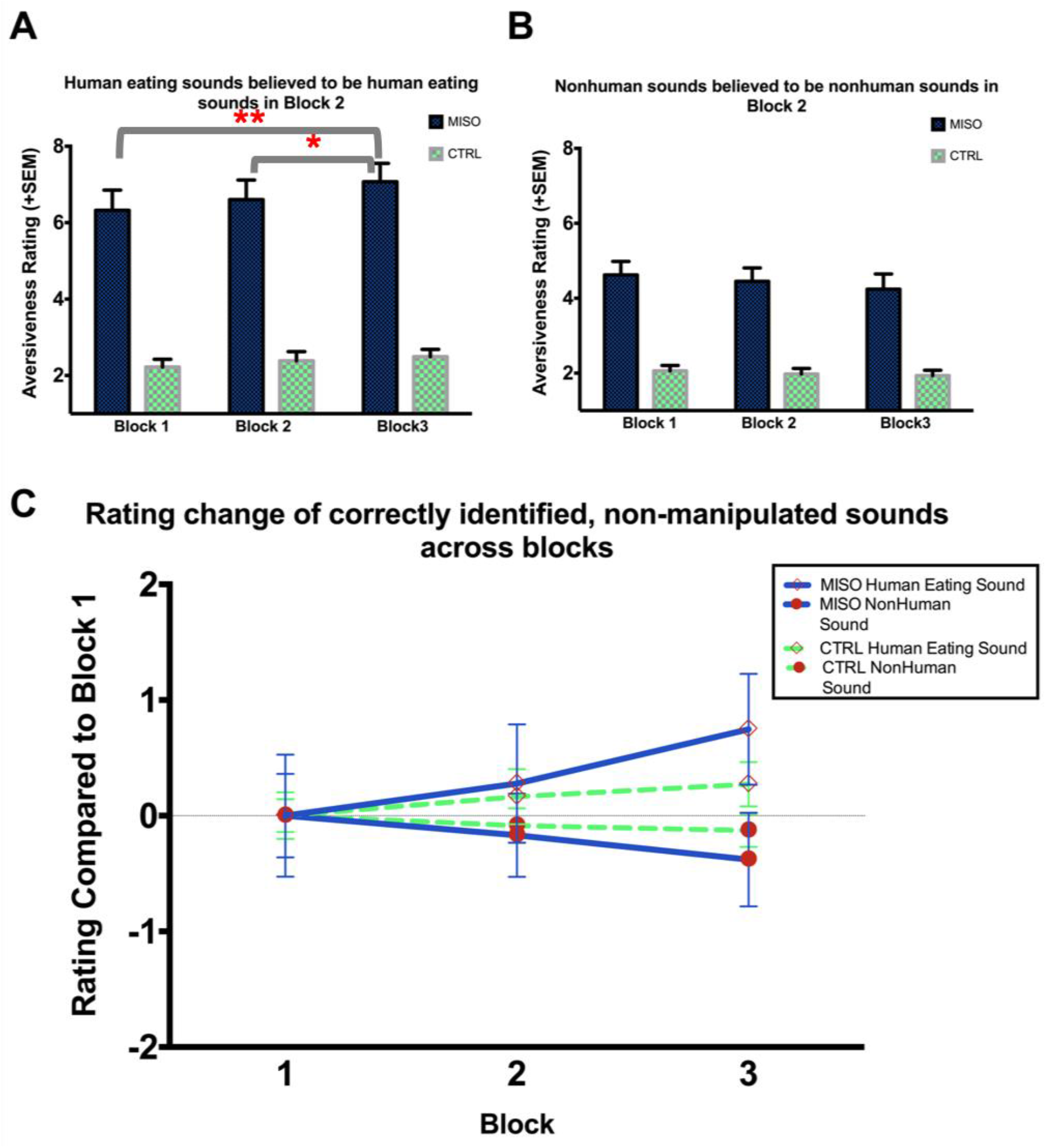
Rating change of human eating sounds and nonhuman sounds (that were correctly identified in all blocks) across blocks. A) Depicts the average misophonic and control aversiveness ratings of human eating sounds that were correctly identified as human eating sounds in blocks 1 and 2. B) Depicts the average misophonic and control aversiveness ratings of nonhuman sounds that were correctly identified as nonhuman sounds in blocks 1 and 2. C) Combines figures 8A and 8B into one graph but instead of displaying average aversiveness ratings, displays the average change in rating of each sound type in each block relative to block 1.

When considering nonhuman sounds (animal eating and non eating sounds) that were correctly identified as nonhuman sounds in blocks 1 and 2, we find that misophonics did not exhibit significant differences in ratings for these sounds between blocks 1 and 2 [p > .05], blocks 2 and 3 [p > .05] or blocks 1 and 3 [p > .05]. Controls also did not exhibit significant differences in ratings for these sounds between blocks 1 and 2 [p > .05], blocks 2 and 3 [p > .05] or blocks 1 and 3 [p > .05] (Fig. 8B).

When comparing human eating sounds that misophonics correctly identified as human eating sounds in blocks 1 and 2 and nonhuman sounds (animal eating and non eating sounds) that misophonics correctly identified as nonhuman sounds in blocks 1 and 2, we find a significant main effect of the between subject factor of Sound Type [F(1,56) = 14.53, p < .001] and a significant interaction between Sound Type and the within subject factor Block [F(2,112) = 3.42, p = .036] (Fig. 8C).

Lastly, when investigating differences in how individual stimuli were rated in block 3 and block 1, we find that misophonic participants had a larger range of difference scores overall than controls. Additionally, although both misophonic and control participants tended to rate human eating sounds as more aversive in block 3 than block 1, and animal eating sounds as less aversive in block 3 than block 1, misophonics demonstrated this to a much greater extent (Fig. 9).

**Figure 9.**
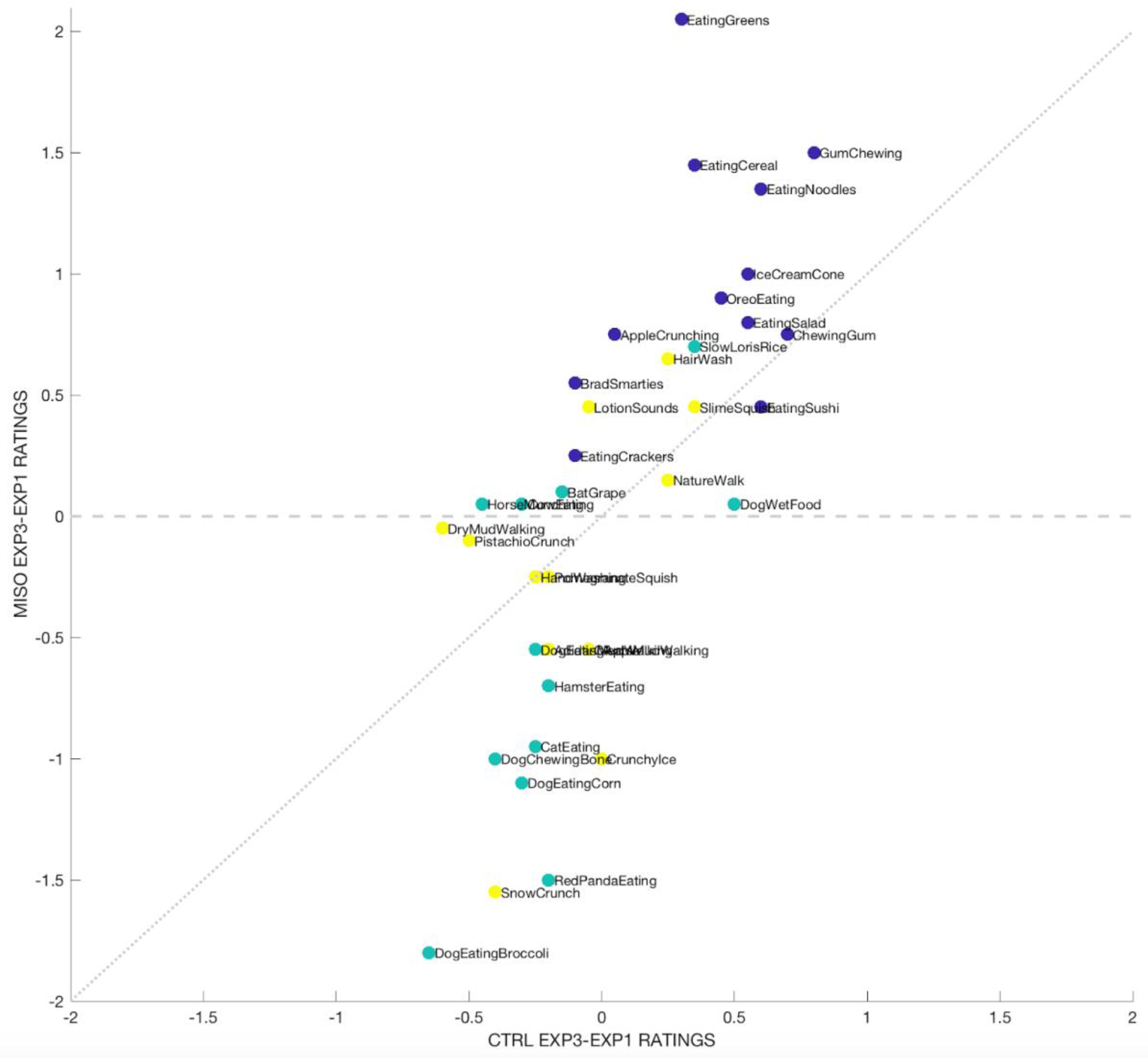
Rating change of individual stimuli between blocks 3 and 1. Shows the difference in rating of each stimulus from when it was encountered in block 1 (audio only) and block 3 (audio + video). Control rating differences are shown on the x axis and misophonic rating differences are shown on the y axis. Purple dots represent human eating sounds, turquoise dots represent animal eating sounds and yellow dots represent non eating sounds.

## DISCUSSION

Overall these results support our main hypothesis that context plays a role in how aversive misophonic participants find certain sounds to be. This is in line with previous reports that suggest that a sound’s source is a crucial factor in determining what is considered a misophonic trigger sound to an individual (Edelstein et al., 2013; Schneider & Arch, 2017). However, the findings from our experiment extend beyond this and show that while sound source is indeed an important factor in the misophonic response, an individual’s *perception* of a sound’s source is enough to influence how they respond to that sound.

Our hypothesis that misophonic participants would find human eating sounds as most aversive when compared with animal eating and non eating sounds was confirmed by within block analyses of ratings and skin conductance of blocks 1, 2 and 3. In block 1, we showed that in the absence of any contextual information (such as text description or video), whether or not a participant correctly guessed a sound’s source (and specifically what they thought the sound was when they didn’t guess correctly), played a role in how aversive they rated that sound to be. Block 2 showed a similar finding, where correct or incorrect text descriptions provided prior to each sound (and whether or not participants believed these descriptions), influenced how aversive participants found those sounds to be. Block 3, which included video of the sound participants were listening to, left no room for interpretation and further solidified the finding that human eating sounds were considered by both misophonic and control participants to be significantly more aversive than animal eating and non eating sounds. Although both groups found human eating sounds to be the most aversive sound category, misophonic individuals always showed much higher aversiveness ratings than controls overall.

In addition to examining how participants responded to these three categories of sounds within blocks, we examined how responses to specific sounds within these categories may change across blocks. In particular, we found that the very same sound could be rated significantly differently from block to block when paired with different contextual information. Specifically, we were interested in the rating change across blocks of human eating and nonhuman sounds (animal eating and non eating sounds were grouped together to form this category) that were identified correctly in block 1, but were believed to be nonhuman sounds and human eating sounds, respectively, when they were heard again in block 2. Indeed, we found that misophonics, but not controls, rated the very same human eating sounds that were correctly identified in block 1, as significantly less aversive when encountered again in block 2 when believed to be nonhuman sounds. When misophonics encountered those same human eating sounds for a third time in block 3, with video, their ratings significantly increased from block 2. Conversely, we found that misophonics, but not controls, rated the very same nonhuman sounds that were correctly identified in block 1, as significantly more aversive when encountered again in block 2 when believed to be human eating sounds. When misophonics encountered those same nonhuman sounds for a third time in block 3, with video, their ratings significantly decreased from block 2.

We were also interested in the rating change across blocks of human eating and nonhuman sounds that were correctly identified as human eating and nonhuman sounds, respectively, in both blocks 1 and 2. While controls did not exhibit significant differences in ratings between blocks for either of these groups of sounds, misophonics did, but only for human eating sounds and not nonhuman sounds. Specifically, misophonics rated human eating sounds as increasingly aversive from blocks 1 to 3. This suggests that for trigger sounds, such as human eating sounds, the more contextual information misophonics are given about what they were listening to, the more aversive the sound becomes.

In terms of future directions, it would be worthwhile to develop a reliable technique to assess physiological markers of misophonia. It should be noted that we collected SCR and electromyography (EMG) data from the participants in this specific study in order to supplement their subjective aversiveness ratings, but unfortunately, due to a number of factors such as a low signal to noise ratio, outdated equipment and the length of the study, not enough of the physiological data ended up being clean enough to be properly analyzed. However, with higher quality recordings, some of the observed main effects from this study would likely produce reliable physiological components.

Ultimately, the findings from this study demonstrate that sound source plays a large role in what are considered to be trigger sounds. The idea that two sounds could sonically sound very similar to each other, but only one might trigger an individual with misophonia, suggested that there is much more that goes into a misophonic trigger than just the sound itself. Through the exclusive use of sonically similar sounds, this study not only showed that human eating sounds were considered to be significantly more aversive than animal eating and non eating sounds to misophonic individuals overall; it also showed that how one interprets these sounds can significantly influence how aversive they believe them to be. The findings from this study show that, depending on the contextual information given, the very same sound could be considered significantly more or less aversive the next time it was encountered. There is already preliminary evidence that cognitive behavioral therapy, which utilizes techniques to help patients reappraise negative thoughts and feelings, may be helpful for individuals with misophonia (Schröder et al., 2017). The fact that there appears to be some degree of cognitive flexibility in terms of reassessing misophonic trigger sounds leads us to believe that there may be successful therapeutic applications of this work in the future.

## Notes

### Competing Interest Statement

The authors have declared no competing interest.

